# Tomato brassinosteroid-signaling kinase Bsk830 is a component of flagellin signaling that regulates pre-invasion immunity

**DOI:** 10.1101/2022.06.01.494411

**Authors:** Guy Sobol, Bharat Bhusan Majhi, Metsada Pasmanik-Chor, Ning Zhang, Holly M. Roberts, Gregory B. Martin, Guido Sessa

**Author notes:** **Correspondence:** Guido Sessa, School of Plant Sciences and Food Security, Tel-Aviv University, 69978 Tel-Aviv, Israel. Department of Chemistry, Biochemistry and Physics, Université du Québec à Trois-Rivières, Trois-Rivières, Quebec G9A 5H7, Canada (Bharat Bhusan Majhi).

## Abstract

Detection of bacterial flagellin by the tomato receptors Flagellin sensing 2 (Fls2) and Fls3 triggers activation of pattern-triggered immunity (PTI). Tomato signaling components associated or downstream of flagellin receptors are largely unknown. We investigated the involvement of tomato brassinosteroid-signaling kinase 830 (Bsk830) in PTI triggered by flagellin perception. Bsk830 localized to the plasma membrane and interacted with Fls2 and Fls3. Consistent with a role in flagellin- induced signaling, CRISPR/Cas9-generated tomato *bsk830* mutants were impaired in ROS accumulation induced by the flagellin-derived flg22 and flgII-28 peptides. In addition, *bsk830* mutants developed larger populations of *Pseudomonas syringae* pv. *tomato* (*Pst*) strain DC3000 than wild-type plants, whereas no differences were observed in plants infected with the flagellin deficient *Pst* DC3000Δ*fliC*. *bsk830* mutants failed to close stomata when infected with *Pst* DC3000 and *Pseudomonas fluorescens,* and were more susceptible to *Pst* DC3000 than wild-type plants when inoculated by dipping, but not by vacuum-infiltration, indicating involvement of Bsk830 in pre-invasion immunity. Analysis of gene expression profiles in *bsk830* mutants detected a reduced number of differentially expressed genes and altered expression of jasmonic acid (JA)-related genes. In support of deregulation of JA response in *bsk830* mutants, these plants were similarly susceptible to *Pst* DC3000 and to the *Pst* DC3118 strain, which is deficient in coronatine production, and more resistant to the necrotrophic fungus *Botrytis cinerea* following PTI activation. These results indicate that tomato Bsk830 is required for a subset of flagellin-triggered PTI responses and support a model in which Bsk830 negatively regulates JA signaling during PTI activation.

## INTRODUCTION

Plants have evolved a complex immune system to confront the wide range of pathogens which inhabit their natural environment. Plant immune responses are activated through recognition of highly conserved microbe-associated molecular patterns (MAMPs) by membrane-localized pattern recognition receptors (PRRs) (DeFalco and Zipfel, 2021), or by detection of pathogen effectors by intracellular nucleotide binding leucine-rich repeat receptors (NLR) (Duxbury et al., 2021). In tomato (*Solanum lycopersicum*), known PRRs include Fls2 (Robatzek et al., 2007) and Fls3 (Hind et al., 2016), which perceive the bacterial flagellin-derived peptides flg22 and flgII-28, respectively, and CORE, which detects the bacterial cold shock protein-derived peptide csp22 (Wang et al., 2016). Flg22 and flgII-28 represent major bacterial MAMPs recognized by tomato (Rosli et al., 2013), and their binding by Fls2 and Fls3 activates signaling pathways that regulate molecular events promoting defense, collectively referred to as pattern-triggered immunity (PTI) (Liang and Zhou, 2018). PTI responses include production of reactive oxygen species (ROS), activation of mitogen-activated protein kinases (MAPKs), transcriptional reprogramming, callose deposition at the cell wall, stomatal closure and activation of hormone signaling (DeFalco and Zipfel, 2021).

Stomata closure is a MAMP-triggered and PRR-mediated defense response, also referred to as stomatal immunity, particularly important against leaf-associated pathogenic bacteria, which gain access to the plant apoplast through these natural openings as well as wounds (Melotto et al., 2017). Dynamics of stomatal immunity against phytopathogenic bacteria have been described for the interaction of Arabidopsis and tomato plants with *Pseudomonas syringae* pv. *tomato* (*Pst*) bacteria (Melotto et al., 2006; Du et al., 2014). Plant infection with *Pst* DC3000 causes stomata closure within 1 h post-inoculation followed by stomata reopening 3-4 h later. Stomatal movement is modulated by ROS accumulation that controls the activity of ion pumps, plasma membrane channels, and transporters (Sierla et al., 2016). In Arabidopsis ROS molecules have been shown to be generated by the NADPH oxidase RBOHD that is activated by phosphorylation upon MAMPs perception by PRRs (Wang and Gou, 2021). MAMP perception also triggers the activation of signaling pathways involving the hormones salicylic acid (SA) and abscisic acid (ABA), which contribute to stomatal closure. Plants defective in SA and ABA synthesis and signaling are unable to induce stomatal closure (Melotto et al., 2017). SA and ABA pathways have been shown to contribute to stomata closure independently and in an interconnected manner (Arnaud and Hwang, 2015).

Stomatal reopening is induced by the pathogen to gain entry into leaves. To promote stomatal reopening, *Pseudomonas syringae* bacteria take advantage of the antagonistic interplay between the plant hormones SA and jasmonic acid (JA) (Thaler et al., 2012). *P. syringae* secretes type III effectors and phytotoxins to enhance JA signaling and repress SA signaling. For example, the HopX1 and HopZ1a effectors induce degradation of JAZ proteins, which are negative regulators of JA signaling, to enhance JA signaling and stomata reopening (Jiang et al., 2013; Gimenez-Ibanez et al., 2014). The *P. syringae* coronatine (COR) phytotoxin is a JA- Ile-mimic molecule that binds the JA receptor COI1 and activates JA-mediated processes (Katsir et al., 2008). Binding of COR to COI1 triggers downstream signaling that induces NAC transcription factors which inhibit SA accumulation and promote stomata reopening (Melotto et al., 2017).

PRRs recruit receptor-like cytoplasmic kinases (RLCKs) to link MAMP perception to downstream signaling. Multiple Arabidopsis RLCKs play a role in plant immunity. For example, BOTRYTIS-INDUCED KINASE1 (BIK1), a member of the RLCK subfamily VII, associates with multiple PRRs and activates downstream signaling components such as the NADPH oxidase RBOHD (Kadota et al., 2014; Li et al., 2014) and several calcium channels (Tian et al., 2019; Thor et al., 2020). Similar to BIK1, additional members of the RLCK subfamily VII contribute to PTI (Rao et al., 2018). RLCKs regulate stomatal immunity and are essential for the initial closure step (Wang and Gou, 2021). For example, BIK1 and PBL27 regulate stomatal closure by promoting activation of ion channels (Zheng et al., 2018; Liu et al., 2019; Thor et al., 2020).

Brassinosteroid-signaling kinases (BSKs) belong to the RLCK subfamily XII and several of them were extensively characterized in Arabidopsis and shown to play a role in brassinosteroid signaling, growth, and response to abiotic stress (Tang et al., 2008; Li et al., 2012b; Sreeramulu et al., 2013; Jia et al., 2019; Ren et al., 2019). Recent investigation identified Arabidopsis BSKs that are involved in plant immunity (Shi et al., 2013; Majhi et al., 2019 and 2021). BSKs participate in various branches of defense signaling, as evident by their interaction with multiple PRRs, and mediation of different PTI responses. For example, BSK1, BSK5, BSK7, and BSK8 associate with the PRR FLS2 and mutations in the corresponding genes compromise a subset of flg22-mediated PTI responses (Shi et al., 2013; Majhi et al., 2019 and 2021). BSK1 was also found to modulate MAPK activation by phosphorylation of the MAPKKK MAPKKK5 and the MAPK MPK15 (Yan et al., 2018; Shi et al., 2022). In addition, while BSK1, BSK7, and BSK8 interact exclusively with FLS2, BSK5 also interacts with EFR and PEPR1 and is required for their signaling (Majhi et al., 2019). In line with a function of BSKs in immune signaling, *bsk1*, *bsk5*, *bsk7* and *bsk8* mutants display enhanced susceptibility to fungal and bacterial pathogens (Shi et al., 2013; Majhi et al., 2019 and 2021). Another family member, BSK3, was shown to interact *in vivo* with multiple components of immune signaling (Xu et al., 2014), but its function in plant immunity is yet unknown.

Tomato BSKs are less characterized, but at least two of the seven family members play a role in plant immunity (Singh et al., 2014; Roberts et al., 2019a). A tomato homolog of BSK7 was found to interact with *Pst* DC3000 effectors and silencing of its homologous genes compromised PTI in *Nicotiana benthamiana* (Singh et al., 2014). Tomato Mai1, homolog of Arabidopsis BSK1, interacts with MAPKKKα, a signaling component required for NLR-mediated immunity (Roberts et al., 2019a). Silencing of Mai1 *N. benthamiana* homologs enhanced susceptibility to *Pst* DC3000 and compromised the hypersensitive response mediated by several *R* genes (Roberts et al., 2019a). Here, we investigated the involvement of tomato BSKs in plant immunity. We provide evidence that Bsk830 physically interacts with flagellin receptors and localizes to the cell plasma membrane. Analysis of two independent loss-of-function mutant lines revealed that Bsk830 is required for stomatal immunity against *Pst* DC3000 and for flagellin-induced ROS production. Analysis of gene expression profiles indicated that loss of *Bsk830* caused an attenuated PTI response and deregulation of JA signaling. Consistent with an altered JA response, loss of *Bsk830* compromised the contribution of the COR toxin to *Pst* DC3000 virulence and reduced susceptibility to a fungal necrotrophic pathogen. Together, our results indicate that Bsk830 modulates a subset of flagellin-induced PTI responses and contributes to regulation of JA signaling during PTI.

## RESULTS

### Tomato Bsk830 interacts with the flagellin receptors Fls2 or Fls3

Recognition of the motility-associated protein flagellin plays a major role in the induction of PTI responses on some tomato accessions (Roberts et al., 2019b). Flagellin contains two MAMPs, flg22 and flgII-28, that are recognized by the PRRs Flagellin sensing 2 (Fls2) and Fls3, respectively (Gómez-Gómez and Boller, 2000; Robatzek et al., 2007; Hind et al., 2016). To investigate the possible involvement of brassinosteroid-signaling kinase (Bsk) proteins in tomato PTI initiated by flagellin perception, we tested interactions of tomato Bsk family members with Fls2 and Fls3 in a yeast two-hybrid system by using Bsk proteins as baits and the kinase domain of Fls2 (Fls2_KD_) and Fls3 (Fls3_KD_) as preys. Bsk830, but none of the other six tomato Bsk family members, interacted with both PRRs (Figure 1A). Next, a split luciferase complementation assay was employed to validate in planta interactions observed in yeast. In these experiments, Fls2_KD_ and Fls3_KD_ were fused to the C-terminal half of the luciferase protein (C-LUC) and co-expressed via *Agrobacterium tumefaciens* in *N. benthamiana* leaves with Bsk830 fused to the N-terminal half of luciferase (N- LUC). As negative controls, C-LUC-PRRs and N-LUC-Bsk830 were co-expressed with N-LUC and C-LUC empty vectors, respectively. Interactions were quantified by measurement of luminescence emitted from leaf discs sampled at 48 h after agro- infiltration. Co-expression of Bsk830 with Fls2_KD_ or Fls3_KD_ resulted in significantly higher luminescence than the negative controls (Figure 1B). Expression in yeast and in planta of all fusion proteins was confirmed by Western blot analysis (Supplemental Figure S1, A and B and Roberts et al., 2019a).

**Figure 1.**
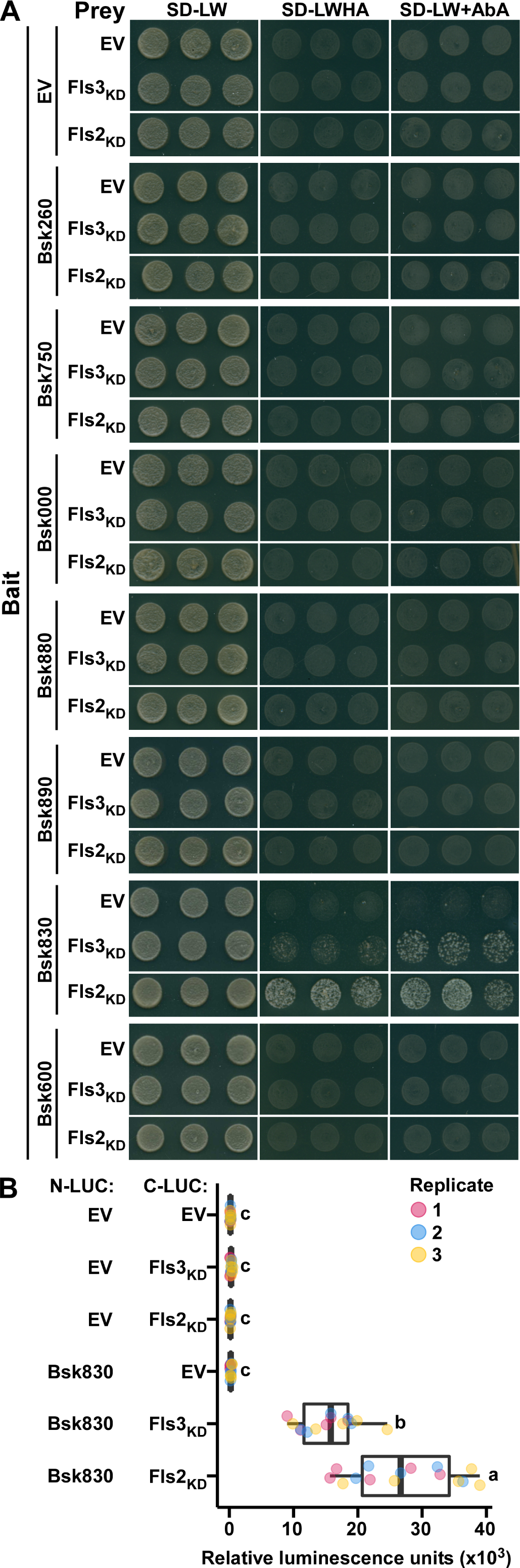
Interaction of Bsk830 with Fls2 and Fls3. A, Yeast cells expressing individual Bsk proteins fused to the GAL4 DNA-binding domain (bait), and the kinase domain of Fls2 (Fls2_KD_) and Fls3 (Fls3_KD_) fused to the GAL4 DNA-activation domain (prey) were grown on synthetically defined medium lacking Leu and Trp (SD–LW), SD–LW lacking His and Ade (SD–LWHA), or SD–LW supplemented with Aureobasidin A (SD–LW+AbA). Empty vectors (EV) were used as negative controls. Nomenclature and accession numbers of tomato Bsk family members are reported in the Materials and Methods section. B, The indicated proteins were fused to C-LUC or N-LUC and co-expressed via *A. tumefaciens* GV2260 in *N. benthamiana* leaves. Luciferase activity was quantified by measuring relative luminescence at 48 h after agro-infiltration. Data from three independent experiments is shown. Circles represent individual data points, and letters represent statistical significance determined by one-way ANOVA and Tukey’s post-hoc test (*P* < 0.05).

To assess the hypothesis that Bsk830 participates in phosphorylation cascade(s) initiated by Fls2 and Fls3, we tested whether Bsk830 is a substrate of Fls2 or Fls3 phosphorylation in an *in vitro* kinase assay. Bsk830 and the cytoplasmic domains of Fls2 (Fls2_CD_) and Fls3 (Fls3_CD_) were fused to the C-terminus of the maltose binding protein (MBP), expressed in *E. coli*, and affinity-purified. MBP-Fls2_CD_ and MBP-Fls3_CD_ were incubated with the MBP-BSK830 fusion in the presence of [γ- ^32^P]ATP in an *in vitro* kinase assay. As previously reported (Roberts et al., 2020), MBP-Fls2_CD_ and MBP-FLS3_CD_ displayed autophosphorylation activity (Supplemental Figure S2). However, phosphorylation of MBP-Bsk830 by MBP-Fls2_CD_ or MBP- Fls3_CD_ was not detected (Supplemental Figure S2), indicating that Bsk830 is not a substrate of Fls2 and Fls3 kinase activity *in vitro* despite its interaction with both PRRs.

### Lipid modifications anchor Bsk830 to the plasma membrane

The Fls2 and Fls3 PRRs are receptor kinases localized to the plasma membrane (PM) (Andolfo et al., 2013; Hind et al., 2016). Based on the interaction of Bsk830 with Fls2 and Fls3, and the presence of putative myristoylation and palmitoylation sites at the Bsk830 N-terminus (Figure 2A), we hypothesized that Bsk830 is anchored to the PM by fatty acids modifications. To examine this possibility, Bsk830 was fused to the N-terminus of YFP and transiently expressed via *A. tumefaciens* in *N. benthamiana* leaves along with the PM marker Flot1b-mCherry (Li et al., 2012a). Bsk830-YFP displayed a similar distribution as Flot1b-mCherry (Figure 2B, upper panels) that was confirmed by quantifying fluorescence detected in the YFP and mCherry channels along a set path (Figure 2B, lower panels). In contrast to Bsk830- YFP, the Bsk830^G2A^-YFP and Bsk830^C(3,11,12)A^-YFP variants, which carry mutations in putative myristoylation and palmitoylation sites, respectively, did not co-localize with Flot1b-mCherry (Figure 2C). These observations suggest that Bsk830 is associated to the PM through myristoylation and palmitoylation modifications.

**Figure 2.**
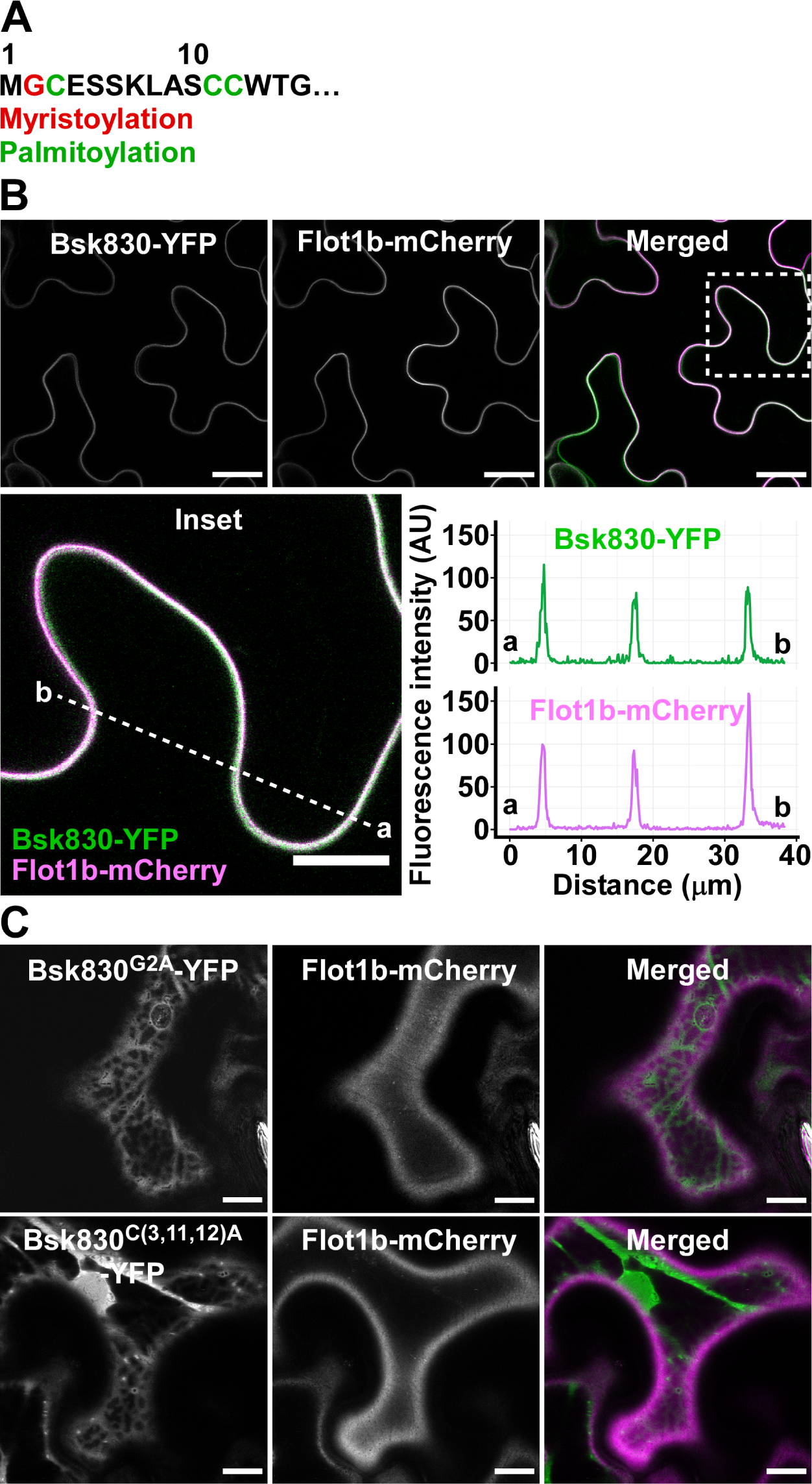
Bsk830 localizes to the plasma membrane. A, Putative myristoylation and palmitoylation sites at the N-terminus of Bsk830. Bsk830-YFP (B), Bsk830^G2A^-YFP (C), or Bsk830^C(3,11,12)A^-YFP (C) fusion proteins were co-expressed via *A. tumefaciens* GV2260 in *N. benthamiana* leaves with the plasma membrane marker Flot1b-mCherry. Fluorescence was monitored in epidermal cells by confocal microscopy at 48 h after agro-infiltration. YFP, mCherry, and merged fluorescence images are shown. The region marked in the merged image by a dotted square is magnified in the inset panel. Fluorescence intensity was measured in the YFP and mCherry channels along the dotted line. Scale bars represents 20 μm, except for the inset image, where it represents 10 μm.

### Mutations in the *Bsk830* gene compromise flagellin-mediated immunity

To investigate the function of Bsk830 in plant immunity, we used CRISPR/Cas9 technology to generate tomato plants with mutations in the *Bsk830* gene. Two independent mutant lines, *bsk830-1* and *bsk830-*2, were generated and allowed to segregate until homozygous genotypes were obtained in the T2 generation. Sequence analysis of the area flanking the gRNA-targeted sequence revealed deletions of 4 bp and 135 bp in the first exon of *Bsk830* in *bsk830-1* and *bsk830-2*, respectively (Supplemental Figure S3). Next, we tested the involvement of Bsk830 in flagellin-induced immunity by examining susceptibility of wild-type and *bsk830* mutant lines to the bacterial pathogen *Pst* DC3000 and its derivative mutant strain *Pst* DC3000Δ*fliC*, which does not form flagella (Kvitko et al., 2009). Plants were inoculated by dipping into suspensions (1x10^7^ CFU/mL) of *Pst* DC3000 and *Pst* DC3000Δ*fliC*, and bacterial populations were determined in leaf tissue at 2 days post-inoculation (dpi). *Pst* DC3000 bacteria displayed a significantly higher growth in *bsk830* mutant lines than in wild-type plants suggesting that Bsk830 is involved in immunity (Figure 3A). In addition, growth of *Pst* DC3000Δ*fliC* was higher than *Pst* DC3000 in wild-type plants, likely because *Pst* DC3000Δ*fliC*, lacking flagellin, is not detected by Fls2 or Fls3. Conversely, growth of *Pst* DC3000 and *Pst* DC3000Δ*fliC* was similar in the *bsk830* mutants, suggesting that in these plants flagellin-induced immunity is impaired. In line with this conclusion, similar bacterial populations were observed in wild-type plants infected with *Pst* DC3000Δ*fliC* and in *bsk830* mutants infected with *Pst* DC3000 indicating that flagellin-mediated immunity was not activated in these interactions: in the first instance because the bacteria did not express flagellin, and in the second instance because the plant was impaired in flagellin signaling.

**Figure 3.**
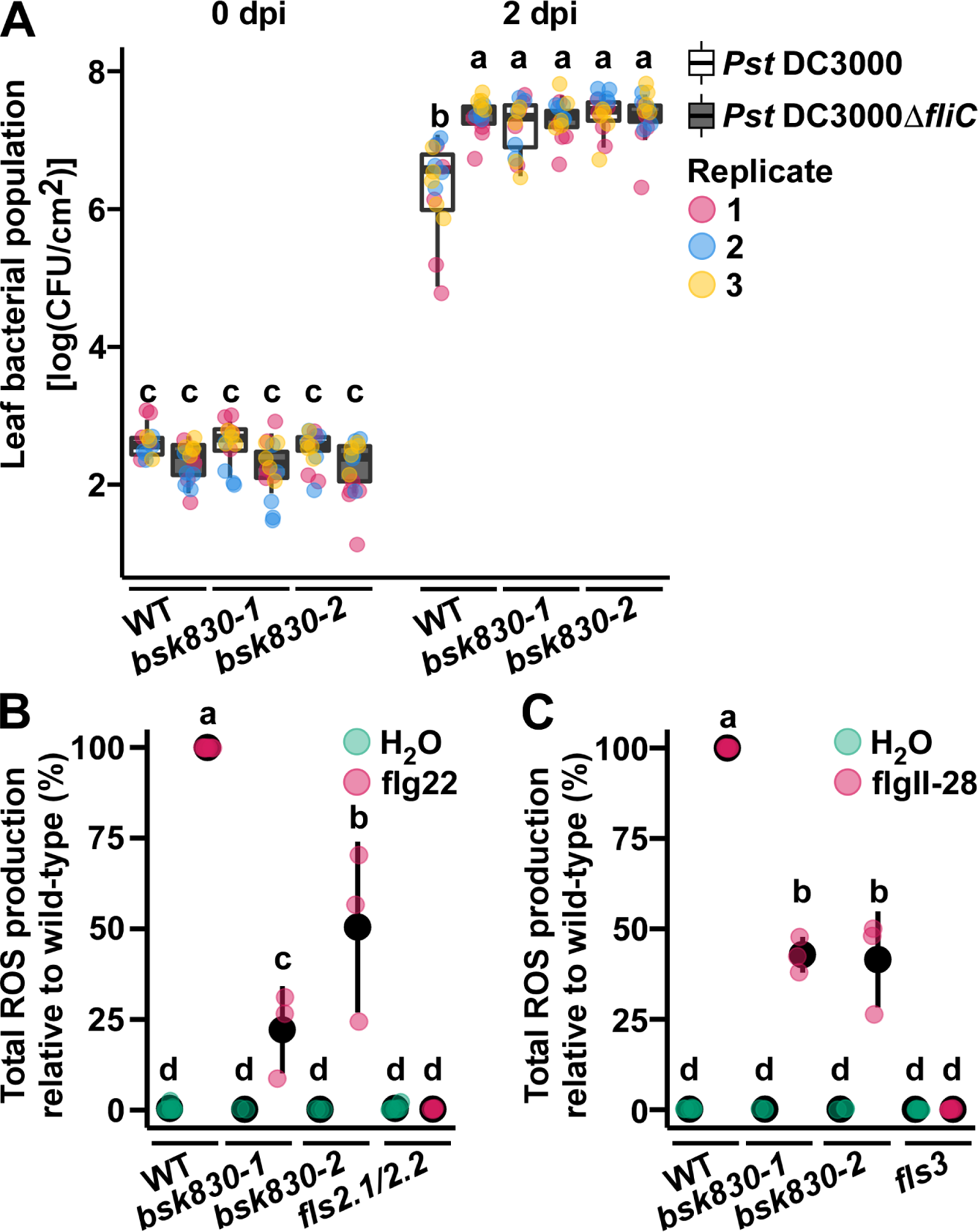
Bsk830 is required for flagellin-mediated immunity. A, Wild-type and *bsk830* mutant plants were inoculated by dipping into a bacterial suspension (10^7^ CFU mL^−1^) of *Pst* DC3000 or *Pst* DC3000Δ*fliC*. Bacterial populations were measured in leaves at 0 and 2 days post-inoculation (dpi). Circles represent individual data points of three biological replicates, and letters represent statistical significance determined by two-way ANOVA and Tukey’s post-hoc test (*P* < 0.05). B and C, ROS production. Leaf discs were treated with 100 nM of flg22, flgII-28, or water. Luminescence was measured for 30 min after flg22 treatment (B) and for 45 min after flgII-28 treatment (C). ROS production was normalized to the ROS amount produced by wild-type plants at its peak. Data are means ± SD of three biological replicates.

To confirm that *bsk830* mutant plants are impaired in flagellin-induced immunity, we examined PTI responses triggered in these plants by the flg22 and flgII-28 MAMPs. Wild-type and mutant plants were treated with flg22 or flgII-28 and monitored for ROS production and MAPK phosphorylation. The *fls2.1/fls2.2* and *fls3* mutants, which are not responsive to flg22 and flgII-28, respectively (Roberts et al., 2020), were used as negative controls. On treatment with flg22 and flgII-28, *bsk830* mutants accumulated lower levels of ROS than wild-type plants (Figure 3, B and C), while MAPK phosphorylation was similarly activated in both genetic backgrounds (Supplemental Figure S4, A and B). As expected, *fls2.1/fls2.2* and *fls3* mutants did not produce ROS (Figure 3, B and C), and were impaired in the activation of MAPK phosphorylation (Supplemental Figure S4, A and B). Together, these results indicate that Bsk830 is required for a subset of flagellin-induced PTI responses.

### *Bsk830* mutant plants are impaired in stomatal immunity

In experiments that revealed enhanced susceptibility of *bsk830* mutants to *Pst* DC3000 infection, plant inoculation was carried out by dipping plants into bacterial suspensions. The use of this inoculation technique left unresolved whether Bsk830 is required for immune responses that counteract the pathogen on the leaf surface during the pre-invasion phase of infection or in the leaf apoplast during the post-invasion phase of infection. To differentiate between these possibilities, wild-type plants and *bsk830* mutants were inoculated with a *Pst* DC3000 suspension by vacuum-infiltration (1x10^5^ CFU/mL), which delivers bacteria directly into the apoplast, or by dipping (1x10^7^ CFU/mL), which requires movement of bacteria through stomata for infection. Bacterial populations were determined in leaf tissues sampled at 0 and 2 dpi. *Pst* DC3000 bacteria displayed a significantly higher growth in *bsk830* mutants than in wild-type plants inoculated by dipping (Figure 4A). However, similar bacterial populations were observed in wild-type and mutants plants inoculated by vacuum-infiltration (Figure 4B), suggesting a role for Bsk830 in pre-invasion immunity. A similar stomatal number index was observed for wild-type and *bsk830* mutant plants (Supplemental Figure S5), excluding a bias due to developmental differences between genetic backgrounds used in these experiments.

**Figure 4.**
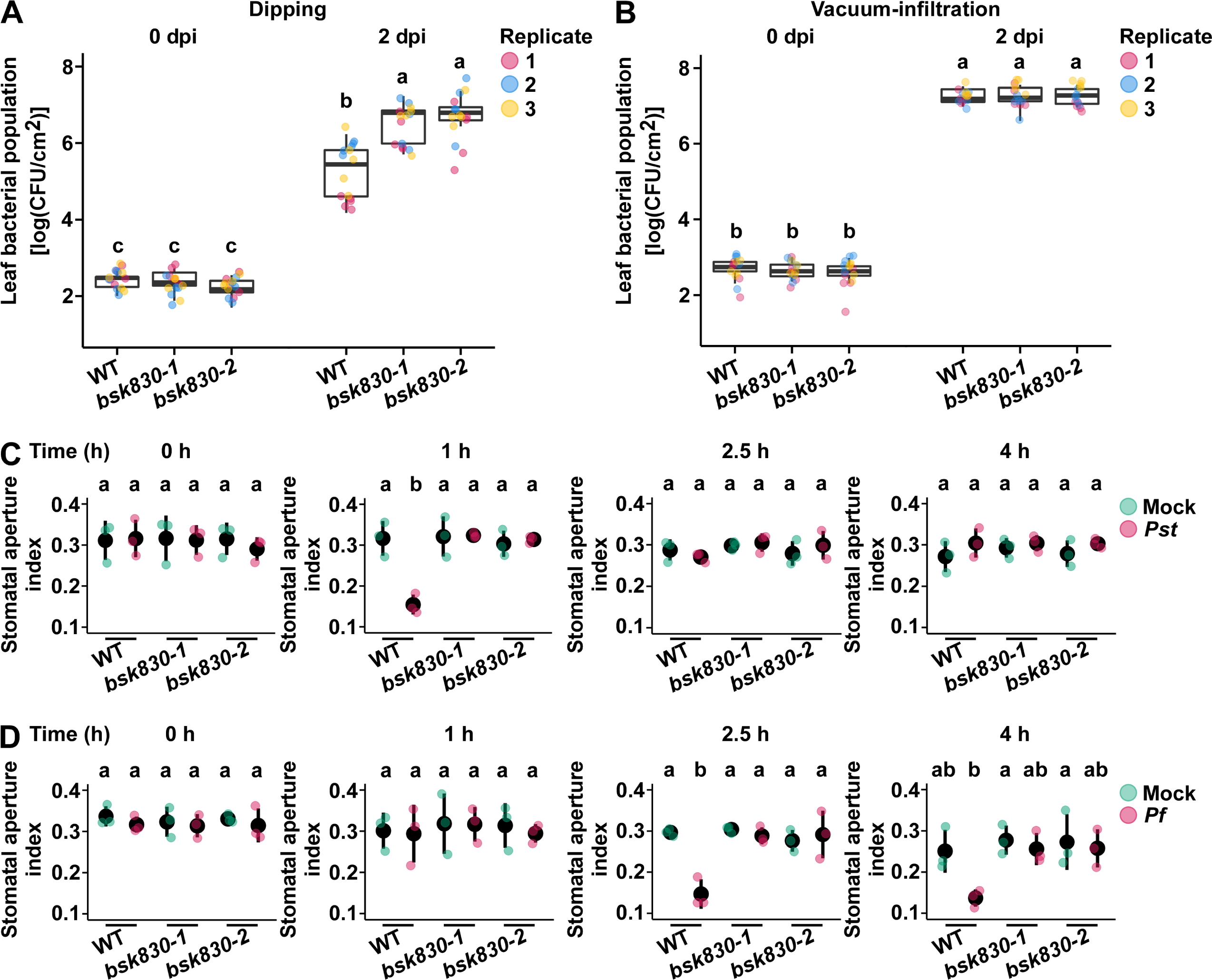
*bsk830* mutant plants are compromised in stomatal immunity. A and B, Wild-type and *bsk830* mutant plants were inoculated with *Pst* DC3000 by dipping (10^7^ CFU mL^−1^) (A) or vacuum-infiltration (10^5^ CFU mL^−1^) (B). Bacterial populations were measured in leaves at 0 and 2 days post-inoculation (dpi). Data from three independent experiments is shown. C and D, Stomatal aperture index (aperture width divided by the stomata length) was determined in leaves after 0, 1, 2.5, and 4 h floating on suspensions (10^8^ CFU mL^−1^) of *Pst* DC3000 (C) and *Pf* A506 (D), or water (mock). Circles represent mean of three biological replicates. Letters represent statistical significance determined by one-way (A and B) or two-way (C and D) ANOVA and Tukey’s post-hoc test (*P* < 0.05).

Because pre-invasion immunity relies on stomatal closure to prevent entrance of bacteria into the leaf apoplast (Melotto et al., 2017), we examined dynamics of stomatal opening/closure in *bsk830* mutants upon infection with *Pst* DC3000 pathogenic bacteria and *Pseudomonas fluorescens* (*Pf*) A506 non-pathogenic bacteria. Both types of bacteria are known to induce stomatal closure early after infection, but only pathogenic bacteria overcome this line of defense at later stages of infection by using virulence factors to reopen stomata (Melotto et al., 2017). Prior to bacterial challeng*e*, leaf pieces of wild-type and *bsk830* mutant plants were floated on stomatal opening buffer under light to ensure stomata opening. After 3 h, leaf pieces were transferred to *Pst* DC3000 or *Pf* bacterial suspension (1x10^8^ CFU/mL), or kept on buffer (mock), and monitored for stomatal apertures during the following 4 h. At 0 h, stomata of both plant genotypes were similarly open, suggesting that loss of function in *Bsk830* does not interfere with light-induced stomatal opening (Figure 4, C and D). On *Pst* DC3000 treatment, stomata of wild-type plants closed at 1 h after infection and reopened at 2.5 h (Figure 4C), while on *Pf* treatment, they closed at 2.5 h and remained closed at 4 h (Figure 4D). In contrast, on both treatments, stomata of the *bsk830* mutant plants remained open for the entire course of the experiment similar to mock-treated plants (Figure 4, C and D). These results indicate that Bsk830 is required for stomatal closure associated with pre-invasion immunity.

### Loss of function in *Bsk830* alters expression of JA- and phenylpropanoid-related genes during PTI

To uncover molecular mechanisms underlying the contribution of Bsk830 to tomato immunity, we compared expression profiles of wild-type and *bsk830* mutant plants during the onset of PTI. Leaves of wild-type and *bsk830-1* lines were inoculated by vacuum-infiltration with a suspension (1x10^8^ CFU/mL) of *Pf* bacteria or a mock solution; samples were collected at 0 and 6 h post-inoculation, and subjected to RNA-seq analysis. In these experiments, we opted for induction of PTI by non-pathogenic rather than by pathogenic bacteria to avoid possible interference of virulence factors. In addition, plants were inoculated by vacuum-infiltration, rather than by dipping, to assure an equal bacterial load in the inoculated leaves despite the defective dynamics of stomatal closure of *bsk830* mutants. A total of 2,146 (852 up-regulated; 1,294 down-regulated) and 1,325 (655 up-regulated; 665 down- regulated) differentially expressed genes (DEGs) were identified in *Pf*- inoculated wild-type and *bsk830* mutant plants, respectively, as compared to mock inoculated plants with filtering parameters of pFDR < 0.05 and a fold change >3 (Figure 5A; Supplemental Table S1). 1,207 DEGs were common to both genotypes, whereas 939 and 118 were unique to wild-type and *bsk830* mutant plants, respectively (Figure 5A). These results suggest that *bsk830* mutants were less responsive to *Pf* inoculation than wild-type plants.

**Figure 5.**
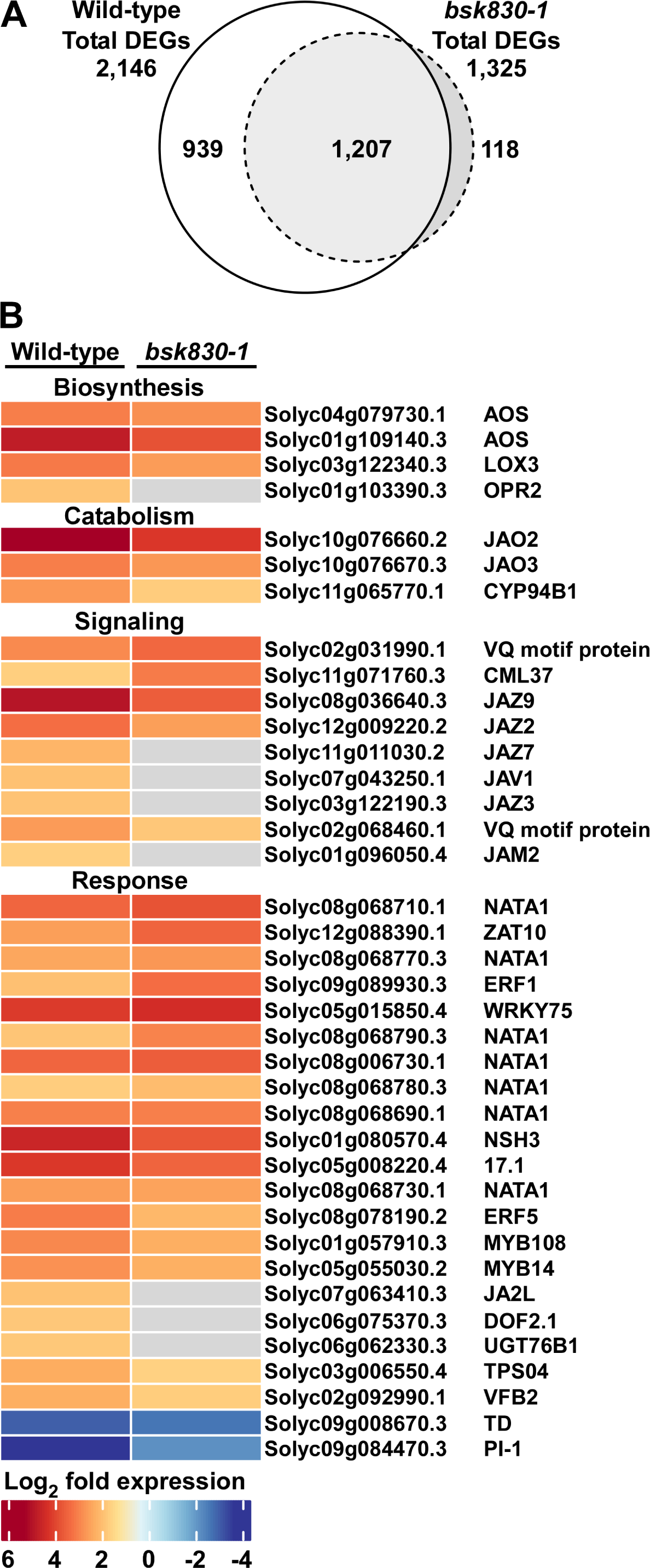
A, Euler diagram representing differentially expressed genes (DEGs) (pFDR < 0.05, fold change > 3, *Pf* vs. mock treatment comparison) in wild-type and *bsk830-1* plants. B, Loss of function in *Bsk830* alters expression of genes involved in JA biosynthesis, catabolism, signaling, and response. Each row represents a single gene accompanied by its Solanaceae Genomics Network accession number. Genes not characterized in tomato are annotated with name of the *Arabidopsis* gene with the highest protein similarity obtained by BLAST. The legend corresponds to relative log_2_ fold change values calculated based on a *Pf* vs. mock treatment comparison performed for either wild-type or the *bsk830* mutant line. Grey rectangles represent genes with no fold change value available.

To identify cellular processes in which Bsk830 is involved, we examined functional categories over-represented among genes that were expressed in wild-type plants, but not in the *bsk830* mutant or vice versa, and genes whose fold-change during PTI differed by at least ±20% between the *bsk830* mutant and the wild-type. Arabidopsis homologs of genes from this pool were subjected to functional enrichment analysis by using the g:Profiler tool (Raudvere et al., 2019) as separate entries based on their up- or down-regulation during PTI. The use of Arabidopsis homologs allowed a more extensive characterization of the gene pool in comparison to the use of tomato gene accessions. Among genes up-regulated during PTI, the predominant functional categories enriched in the *bsk830* mutant were related to JA signaling and response, and to metabolism of phenylpropanoids (Table 1). In *bsk830* mutant, expression of genes involved in JA biosynthesis (e.g., AOS, LOX3, OPR2; Wasternack and Song, 2016), catabolism (e.g., JAO2, JAO3, CYP94B1; Smirnova et al., 2017), and negative regulation of JA signaling (e.g., JAZ2, JAZ3, JAZ7, JAZ9, JAM2; Sasaki-Sekimoto et al., 2013) was reduced as compared to wild-type plants (Figure 5B, Supplemental Table S1). In addition, the transcript abundance of genes encoding various JA response factors (e.g., NATA1, TD, PI-I, ERF1, ERF5, JA2L; Du et al., 2017) was reduced or increased (Figure 5B, Supplemental Table S1). Genes related to phenylpropanoid metabolism displayed increased expression in the *bsk830* mutant and included homologs of phenylalanine ammonia-lyase (PAL) as well as enzymes acting downstream to PALs and involved in lignin biosynthesis (Vanholme et al., 2019) (Supplemental Table S1). Among genes down-regulated during PTI, the predominant categories enriched in the *bsk830* mutant were mainly related to photosynthesis (Table 1). Expression of photosynthesis-related genes was more elevated in the *bsk830* mutant than in wild-type plants (Supplemental Table S1), suggesting a less extensive reallocation of resources from general metabolism to defense that is usually observed in plants activating immune responses (Attaran et al., 2014). Together, these results suggest that loss of *Bsk830* results in a weaker PTI response and altered regulation of JA signaling and response.

**Table 1.** Functional categories overrepresented in DEGs of *Pf* treated *bsk830* plants.

| **Up-regulated genes** |  |  |
| --- | --- | --- |
| **Term name** | **Term ID** | **-log<sub>10</sub>(p<sub>adj</sub>)** |
| response to chitin | GO:0010200 | 5.53039 |
| regulation of jasmonic acid mediated signaling pathway | GO:2000022 | 5.42490 |
| cellular response to fatty acid | GO:0071398 | 4.02940 |
| jasmonic acid mediated signaling pathway | GO:0009867 | 3.47824 |
| cellular response to jasmonic acid stimulus | GO:0071395 | 3.37033 |
| lignin metabolic process | GO:0009808 | 3.19087 |
| regulation of defense response to insect | GO:2000068 | 3.18180 |
| phenylpropanoid metabolic process | GO:0009698 | 2.80966 |
| phenylpropanoid biosynthetic process | GO:0009699 | 2.19778 |
| regulation of anion channel activity | GO:0010359 | 2.09635 |
| **Down-regulated genes** |  |  |
| **Term name** | **Term ID** | **-log<sub>10</sub>(p<sub>adj</sub>)** |
| photosynthesis, light reaction | GO:0019684 | 44.88674 |
| photosynthetic electron transport chain | GO:0009767 | 21.43202 |
| photosynthesis, light harvesting | GO:0009765 | 17.15130 |
| photosynthesis, light harvesting in photosystem I | GO:0009768 | 14.07682 |
| electron transport chain | GO:0022900 | 14.04188 |
| photosynthetic electron transport in photosystem I | GO:0009773 | 13.60706 |
| NAD(P)H dehydrogenase complex assembly | GO:0010275 | 11.56992 |
| photosystem II assembly | GO:0010207 | 7.293742 |
| response to high light intensity | GO:0009644 | 5.682999 |
| glucose metabolic process | GO:0006006 | 5.284815 |

### Mutation of *Bsk830* affects JA-mediated phenotypes

To assess the hypothesis that loss of *Bsk830* causes altered regulation of the JA response during the onset of PTI, we examined the contribution of the COR toxin to *Pst* DC3000 virulence in wild-type and *bsk830* mutant plants. COR is a hormone mimic, which closely resembles JA-Ile (Katsir et al., 2008), and it has been shown to activate JA signaling and promote stomatal opening (Melotto et al., 2017). Leaves of wild-type, *bsk830-1* and *bsk830-2* lines were inoculated by dipping in bacterial suspensions (1x10^7^ CFU/mL) of *Pst* DC3000 (able to synthesize COR; COR+) or *Pst* DC3118 (unable to synthesize COR; COR-), and bacterial populations were determined in leaves at 0 and 2 dpi. In wild-type plants, *Pst* DC3000 (COR+) bacteria displayed a significantly higher growth than *Pst* DC3118 (COR-) (Figure 6A) indicating that COR promotes bacterial virulence, as previously reported (Zheng et al., 2012; Du et al., 2014; Gimenez-Ibanez et al., 2017). Conversely, in *bsk830* mutant plants, *Pst* DC3000 (COR+) and *Pst* DC3118 (COR-) displayed a similar growth indicating that mutation of *Bsk830* compromises the contribution of COR to bacterial virulence and suggesting that, similar to COR, a mutation in *Bsk830* deregulates the JA response.

**Figure 6.**
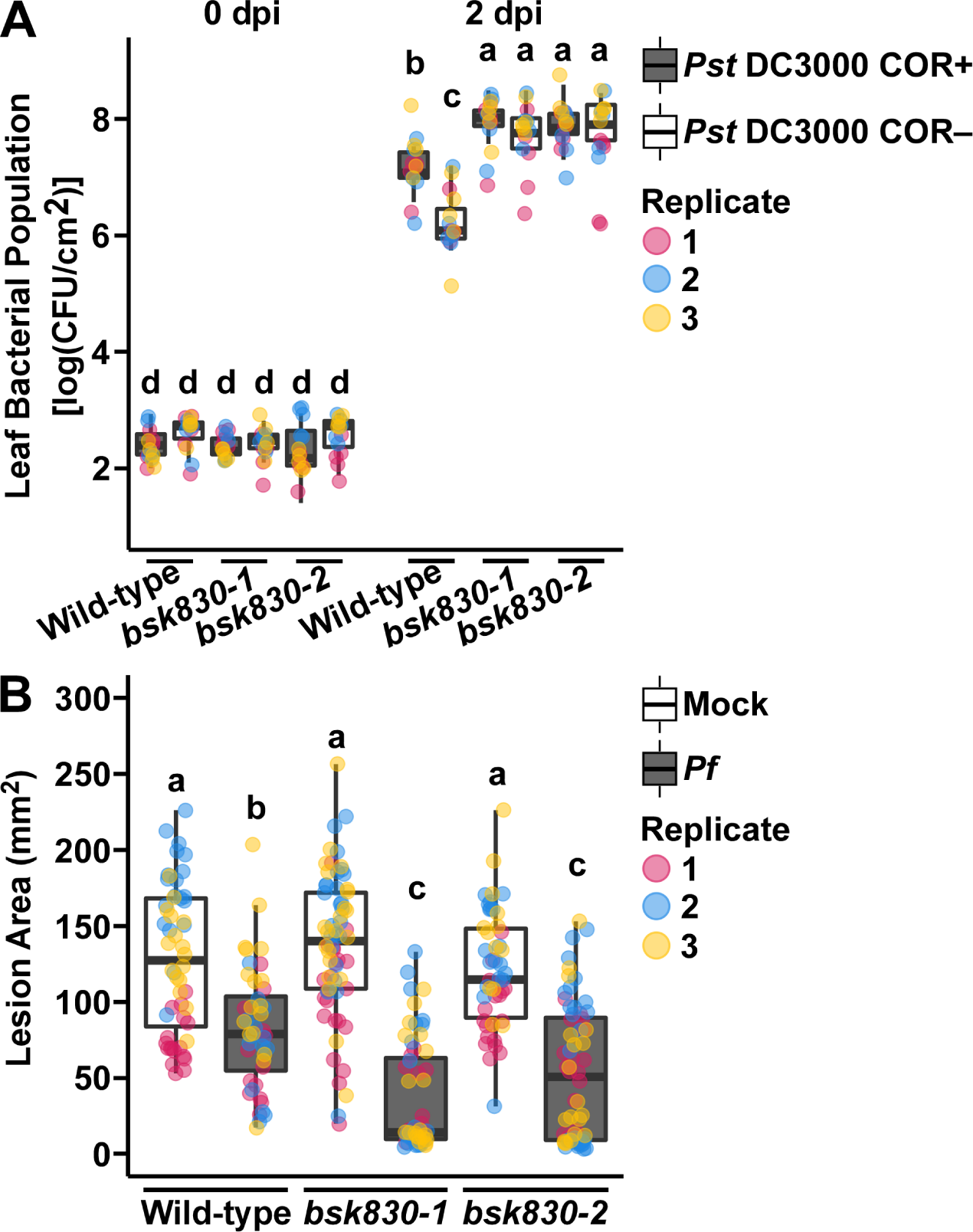
A, Plants of the indicated genotypes were inoculated by dipping with a 10^7^ CFU mL^−1^ bacterial suspension of *Pst* DC3000 (COR+) or *Pst* DC3118 (COR–). Bacterial populations in leaves were measured at 0 and 2 dpi. B, Plants of the indicated genotypes were vacuum-infiltrated with a *Pf* A506 suspension (10^8^ CFU mL^−1^). After 24 h, plants were inoculated by placing a droplet of a suspension carrying *Botrytis cinerea* spores (2 × 10^5^ conidia mL^−1^). Lesion area was measured at 3 dpi. In (A and B) data from three independent experiments is shown. Letters represent statistical significance determined by two-way ANOVA and Tukey’s post- hoc test (*P* < 0.05).

Next, we examined the effect of loss of function in *Bsk830* on susceptibility to the necrotrophic fungal pathogen *Botrytis cinerea*, which is known to be mediated by JA (Zhang et al., 2017). Wild-type and *bsk830* mutant plants were pretreated with a mock solution or with a *Pf* bacterial suspension (1x10^8^ CFU/mL) to activate PTI, and 24 h later plants were infected by placing on the leaves a droplet of a *B. cinerea* spore suspension (2 × 10^5^ conidia mL^−1^). Lesion diameter in infected leaves was measured at 3 dpi (Figure 6B). Wild-type and *bsk830* mutant plants were equally susceptible to *B. cinerea* when mock treated. However, following the *Pf* treatment, symptoms developed more slowly and lesions were significantly smaller in leaves of *bsk830* mutant plants than in wild-type plants. These results support a model in which the JA response is deregulated during the onset of PTI in *bsk830* mutant plants.

## DISCUSSION

We identified the tomato RLCK Bsk830 as a component of signaling pathways originated from perception of bacterial flagellin by the Fls2 and Fls3 PRRs. Initial indication for the involvement of Bsk830 in plant immunity was its interaction with the flagellin receptors Fls2 and Fls3 that was observed in yeast and then validated in planta. Subsequent analysis of *bsk830* mutant plants revealed that Bsk830 is required for pre-invasion immunity by mediating a subset of PTI responses including ROS accumulation and stomatal closure possibly through negative regulation of JA signaling. BSK family members appear to play a role in flagellin-induced signaling in different plant species: in tomato, Bsk830 (but not other Bsk family members) is recruited by two flagellin receptors and is required for flagellin-induced immunity, while in Arabidopsis multiple BSKs (BSK1, BSK5, BSK7, BSK8) interact with the FLS2 PRR and play a role in flg22-induced PTI (Shi et al., 2013; Majhi et al., 2019 and 2021). It remains to be determined whether Bsk830 participates only in signaling pathways activated by flagellin-derived MAMPs, similar to its closest Arabidopsis homologs BSK7 and BSK8 (required for PTI responses triggered by flg22, but not by elf18 and pep1 [Majhi et al., 2021]), or if it is involved in signaling events activated by multiple MAMPs, as observed for Arabidopsis BSK5 (Majhi et al., 2019).

Similar to other BSK family members (Majhi et al., 2019 and 2021; Ren et al., 2019; Roberts et al., 2019a; Su et al., 2021), Bsk830 localizes to the plasma membrane, where it may interact with PRRs and associated components of immune complexes. N-terminal sites predicted to mediate myristoylation and palmitoylation modifications were essential to its plasma membrane localization, common mechanisms used by BSKs for plasma membrane anchoring (Majhi et al., 2019 and 2021; Ren et al., 2019; Roberts et al., 2019a; Su et al., 2022) or maintenance of protein stability, as demonstrated for BSK1 (Su et al., 2022). However, the output of the interaction of Bsk830 with Fls2 and Fls3 is yet unknown. It is unlikely that Bsk830 is activated by Fls2 and/or Fls3 phosphorylation, because Fls2 and Fls3 were not able to phosphorylate Bsk830 *in vitro* despite detection of their autophosphorylation activity. It is possible that additional molecules, which participate *in vivo* in Bsk830 phosphorylation by Fls2 and Fls3, are missing *in vitro*. Alternatively, Bsk830 might function as a scaffolding protein that mediates signal transduction by bringing signaling components in proximity, as it has been suggested for other BSKs (Sreeramulu et al., 2013; Ren et al., 2019; Majhi et al., 2019 and 2021). The latter hypothesis is supported by the evidence that Bsk830 autophosphorylation was not detected under the conditions used in our *in vitro* experiments.

Phenotypic analysis of *bsk830* mutant lines revealed that Bsk830 is required for a subset of PTI responses induced by flagellin-derived MAMPs, including ROS production, but not MAPK phosphorylation. This result, together with previous observations that Arabidopsis *bsk1*, *bsk5*, *bsk7* and *bsk8* mutant lines are impaired in flg22-induced ROS production (Shi et al., 2013; Majhi et al., 2019 and 2021), confirms a central role of BSK family members in signaling pathway(s) that link flg22 sensing to ROS accumulation. Conversely, it is less likely that BSK family members are involved in signaling of flg22-induced MAP kinase activation. In our experiments, a mutation in *Bsk830* did not alter MAP kinase activation induced by flg22 treatment in tomato plants. Similarly, Arabidopsis lines carrying different combinations of mutations in *BSK* genes, including up to seven *BSK*s, displayed a similar MAPK activation as wild-type plants following flg22 challenge (Majhi et al., 2021). However, it is still possible that BSKs participate in MAP kinase activation induced by other MAMPs or pathogen effectors. In support of this hypothesis, BSK1 was shown to phosphorylate *in vitro* MAPKKK5 and MPK15, and required for disease resistance mediated by these MAPKKKs against virulent and avirulent *Pst* DC3000 strains and powdery mildew fungi, respectively (Yan et al., 2018; Shi et al., 2022).

We provide evidence that Bsk830 is involved in pre-invasion immunity, as *bsk830* mutants failed to close stomata when inoculated with pathogenic and non-pathogenic bacterial strains, and displayed enhanced susceptibility to *Pst* DC3000 when inoculated by dipping, but not by vacuum-infiltration. Stomata closure is a PTI response that prevents bacteria from entering the plant apoplast and is activated by detection of MAMPs by PRRs (Melotto et al., 2017). For example, Arabidopsis plants mutated in FLS2 or in both PEPR1 and PEPR2 PRRs fail to close stomata when challenged with the respective MAMPs, and are more susceptible to *Pst* DC3000 when dip-inoculated, but not when syringe-infiltrated (Zipfel et al., 2004; Melotto et al., 2006; Zheng et al., 2018). In Arabidopsis, the RLCKs BIK1 and PBL27 act downstream of PRRs in signaling pathways that lead to stomata closure (Wang and Gou, 2021). BIK1 transduces the signal generated by recognition of flg22 and pep1 by their respective PRRs and activates anion channels that cause stomata closure (Kadota et al., 2014; Li et al., 2014; Guzel Deger et al., 2015; Zheng et al., 2018; Thor et al., 2020). Similarly, PBL27, interacts with the chitin receptor CERK1 and upon chitin elicitation activates anion channels that cause stomata closure (Liu et al., 2019). RLCKs of the BSK subfamily were not examined in the context of stomatal movement, with the exception of the observation that *BSK5* mutant plants are hypersensitive to ABA in stomatal closure (Li et al., 2012b).

Comparison of gene expression profiles of *bsk830* mutant and wild-type tomato plants during the activation of PTI allowed us to formulate hypotheses about the molecular mechanisms of Bsk830 to regulate PTI responses. A prominent difference between *bsk830* mutants and wild-type plants was differential expression of genes related to JA biosynthesis, catabolism, signaling, and response. JA regulates stomatal aperture by promoting their opening, as opposed to SA that promotes stomata closure in response to bacterial pathogens in Arabidopsis and tomato plants (Lee et al., 2013; Melotto et al., 2017; Guzman et al., 2020). Pathogens manipulate stomata aperture by secretion of JA mimicking molecules or effectors (Melotto et al., 2017). For example, *Pst* DC3000 secretes COR, a hormone mimic that closely resembles JA-Ile, that promotes degradation of JAZ proteins which negatively regulate transcription of JA-related genes and signaling (Zhang et al., 2017).

Activation of the JA pathway leads to an inhibitory effect on accumulation of salicylic acid, which in turn promotes stomatal reopening (Melotto et al., 2017). Our expression profiles data indicates a release of negative regulation of JA signaling in the *bsk830* mutants compared to wild-type plants. This was confirmed by the lack of COR contribution to *Pst* DC3000 virulence in *bsk830* mutants, and their increased resistance to a fungal necrotrophic pathogen following PTI activation (Zhang et al., 2017). We therefore propose a model in which Bsk830 negatively regulates JA signaling and response that promote stomatal closure and ROS production (Yi et al., 2014; Toum et al., 2016). This is reminiscent of other regulators of plant immunity, such as FERONIA, which destabilizes MYC2, a regulator of JA signaling, to inhibit COR-induced signaling and promote disease resistance (Guo et al., 2018), and LINC1, which negatively regulates transcription of JA-related genes to enhance PTI (Jarad et al., 2020). In conclusion, our data reveal an important role for tomato Bsk830 in pre-invasion immunity initiated by flagellin perception and in regulation of JA signaling and response during PTI.

## MATERIALS AND METHODS

### Plant materials and growth

Plant cultivars used were: *Nicotiana benthamiana* (Goodin et al., 2008), and tomato (*Solanum lycopersicum*) Hawaii 7981 (Wang et al., 2011). *N. benthamiana* plants were grown in a phytochamber at 25°C in long-day conditions (16 h/8 h, light/dark). Tomato plants were grown in a greenhouse with temperatures fluctuating between 25°C to 30°C under natural light conditions.

### Strains and growth conditions of bacteria, fungi, and yeast

The strains used were: *Escherichia coli* DH5α (Invitrogen) and Rosetta (MERCK), *Pseudomonas syringae* pv. *tomato* (*Pst*) DC3000 (Guo et al., 2009), *Pst* DC3000Δ*fliC* (Kvitko et al., 2009), *Pst* DC3118 (Ma et al., 1991), *Pseudomonas fluorescens* A506 (Wilson et al., 2002), *Agrobacterium tumefaciens* GV2260 (Deblaere et al., 1985) and LBA4404 (Ooms et al., 1982), *Botrytis cinerea* B05.10 (Ma et al., 2017), and yeast (*Saccharomyces cerevisiae*) Y2HGold (Clontech). Bacterial, fungal, and yeast strains were grown with the appropriate antibiotics as follows: *E. coli* in Lysogeny broth (LB) medium at 37°C; *Pst*, *Pf*, and *A. tumefaciens* in LB at 30°C; *B. cinerea* in potato dextrose broth at 20°C; yeast in synthetically defined medium (6.7 g/L yeast nitrogen base without amino acids, 1.4 g/L amino acid drop-out mix) supplemented with 2 % (w/v) glucose at 30°C.

### Pathogenicity assays

Tomato plants were inoculated by vacuum-infiltration with bacterial suspensions of 1x10^5^ CFU/mL in 10 mM MgCl_2_ and 0.008% (v/v) Silwet L-77 (apart from *Pf* A506, which was inoculated at a concentration of 1x10^8^ CFU/mL), or by dipping into bacterial suspensions of 1x10^7^ CFU/mL in 10 mM MgCl_2_ and 0.04% Silwet L-77. Plants inoculated by dipping were placed in a sealed transparent box to maintain humidity. Leaflets were with 3% (w/v) bleach, rinsed with water, and dried. Five samples of four leaf discs (1 cm diameter) were taken from three plants at 2 h after inoculation (day 0) and 2 days later, and homogenized in 1 mL of 10 mM MgCl_2_ to determine bacterial populations via serial dilution plating. For tomato inoculation with *Botrytis cinerea,* droplets (7 µL) of 0.5X potato dextrose broth containing 2x10^5^ spores/mL were deposited on the leaf surface. The area of disease lesions was measured three days after inoculation.

### Peptide elicitors

Peptides flg22 (QRLSTGSRINSAKDDAAGLQIA; Krol et al., 2010) and flgII-28 (ESTNILQRMRELAVQSRNDSNSSTDRDA; Clarke et al., 2013) were purchased from Integrated DNA Technologies, Inc., dissolved in water to a 5 mM stock solution, and diluted to the working concentration.

### *A. tumefaciens*-mediated transient expression

Overnight cultures of *A. tumefaciens* were pelleted, washed three times with 10 mM MgCl_2_, resuspended in induction medium (10 mM MgCl_2_, 10 mM MES [pH 5.6], and 200 μM acetosyringone), and incubated with shaking for 3-4 h at 20°C. Cultures were diluted to OD_600_=0.2 and infiltrated into leaves of *N. benthamiana* plants using a needleless syringe.

### Generation of CRISPR/Cas9-mediated knockout lines

To generate tomato lines with mutations in the *Bsk830* gene, guide RNAs (gRNA1: GATTCTGAGCCTCGTGAATG; gRNA2: GTTTAACAGCAACCGGCCTC) targeting the first exon of *Bsk830* were designed using the tomato genome version SL2.5 (Tomato Genome Consortium, 2012) and the Geneious R11 software (https://www.geneious.com). Each gRNA was cloned into a Cas9-expressing binary vector (p201N:Cas9; Jacobs et al., 2015) by Gibson assembly (Jacobs et al., 2017) and transformed into *A. tumefaciens* LBA4404. The obtained strains were pooled and used to transform Hawaii 7981 tomato plants at the Boyce Thompson Institute transformation facility (Frary and Van Eck, 2005). To determine the mutation type, genomic DNA was extracted from transgenic leaves using a modified CTAB method (Murray and Thompson, 1980). Genomic regions flanking the target site of the *Bsk830* gene were amplified with primers 15-16 (Supplemental Table S2) and sequenced.

### Yeast two-hybrid assay

A GAL4 two-hybrid system was used to analyze protein-protein interactions in yeast. pGBKT7 vectors (bait) carrying tomato Bsk family members fused to the GAL4 DNA binding domain were as described (Roberts et al., 2019a). Gene fragments encoding the kinase domains of Fls2 (Fls2_KD_; amino acids 867 to 1,169) or Fls3 (Fls3_KD_; amino acids 854 to 1,140) were PCR-amplified from tomato cDNA using primers 3-4 (Fls2_KD_) or 1-2 (Fls3_KD_) (Supplemental Table S2) and fused to the GAL4 activation domain in the pGADT7RecM vector (prey). Interactions were tested by placing droplets of yeast (10 μl) carrying bait and prey vectors on SD medium lacking Leu and Trp (SD−LW), SD−LW lacking His and adenine (SD-LWHA), or SD−LW supplemented with 25 μg/ml Aureobasidin A (SD–LW+AbA).

### Split luciferase complementation assay

Gene fragments encoding Fls2_KD_ and Fls3_KD_ were PCR-amplified from tomato cDNA using primers 7-8 (Fls2_KD_) or 5-6 (Fls3_KD_) (Supplemental Table S2), and cloned in frame to the C-terminal fragment of firefly luciferase in the binary vector pCAMBIA1300:C-LUC (Chen et al., 2008). Bsk830 was cloned in frame to the N-terminal fragment of firefly luciferase in the binary vector pCAMBIA1300:N-LUC (Chen et al., 2008), as described (Roberts et al., 2019a). The obtained vectors were transformed into *A. tumefaciens* GV2260 and co-expressed in *N. benthamiana* leaves. Split luciferase complementation assays were performed as described (Majhi et al., 2019).

### Subcellular localization

Gene fragments encoding *Bsk830*, *Bsk830*^G2A^, and *Bsk830*^C(3,11,12)A^ were PCR- amplified from tomato cDNA using primers 9-10, 11-10, and 12-10, respectively (Supplemental Table S2). Amplified fragments were fused upstream of the coding region of the yellow fluorescence protein (YFP) in the pBTEX binary vector driven by the CaMV 35S promoter (Frederick et al., 1998; Popov et al., 2016). Fusion proteins were co-expressed with the plasma membrane marker Flot1b-mCherry (Li et al., 2012a) via *A. tumefaciens* GV2260 in *N. benthamiana* leaves and their localization was visualized by a Zeiss LSM780 confocal laser scanning microscope (Zeiss). YFP was excited with an argon laser at 514 nm, while mCherry and mRFP were excited with a DPSS561-10 laser at 561 nm. Emission was detected between 518 and 583 nm for YFP, and between 579 and 650 nm for mCherry and mRFP. Images were processed with the image processing package Fiji (https://fiji.sc/).

### Expression of MBP fusion proteins in *E. coli* and *in vitro* kinase assay

A gene fragment encoding Bsk830 was PCR-amplified from tomato cDNA using primers 13-14 (Supplemental Table S2), and cloned into the pMAL-c2x vector (New England Biolabs). Fls2_CD_ (amino acids 841 to 1,169), and Fls3_CD_ (amino acids 824 to 1,140) MBP fusions in the pMAL-c2x vector were prepared as described (Roberts et al., 2020). Proteins were expressed in the Rosetta *E. coli* strain and purified (Majhi et al., 2019). MBP fusion proteins were incubated at 25°C for 1 h in a kinase assay solution containing 50 mM Tris-HCl (pH 7), 1 mM DTT, 10 mM MgCl_2_, 10 mM MnCl_2_, 20 µM ATP, and 10 µCi [γ-^32^P]ATP (3,000 Ci mmol^−1^; Perkin-Elmer). Reactions were stopped by addition of the SDS sample buffer. Half of the reaction was fractionated by SDS-PAGE and stained with Coomassie Blue. The second half was fractionated by SDS-PAGE, blotted onto a PVDF membrane and exposed to autoradiography.

### Protein extraction

Protein extraction from yeast and leaves was performed as described by Salomon and Sessa (2010) and Majhi et al. (2019), respectively.

### ROS production assay

Leaf discs were placed in 200 μL of water overnight with the adaxial side up in 96- well plates. The next day, the water was replaced with a solution containing 100 nM flg22 or flgII-28, 34 μg mL^−1^ luminol (Sigma-Aldrich), and 20 μg mL^−1^ horseradish peroxidase (Sigma-Aldrich). Luminescence was measured with a Veritas Microplate Luminometer (Turnerbiosystems Veritas) for 30 min (flg22) or 45 min (flgII-28) at 2.5- or 3-min intervals, with a reading time of 2 sec per well.

### MAPK phosphorylation assay

Leaf discs (∼60 mg) were floated overnight in 10 mL water in 6-well-plates and then treated with 1 µM flg22, 1 µM flgII-28, or water. Total proteins were extracted in 150 µL extraction buffer (50 mM Tris-HCl [pH 7.5], 200 mM NaCl, 1 mM EDTA, 10 mM NaF, 1 mM Na_2_MoO_4_, 2 mM Na_3_VO_4_, 10% [v/v] glycerol, 2 mM DTT, and 1 mM PMSF). Proteins were fractionated by SDS-PAGE, transferred onto PDVF membranes, and immunoblotted with rabbit anti-pMAPK antibodies (α-pMAPK; Cell Signaling Technology).

### Measurements of stomatal aperture and number

Leaf pieces were cut from tomato leaflets and floated on stomatal buffer (25 mM MES-KOH [pH 6.15] and 10 mM KCl) for 3 h under light to allow stomata to fully open (Melotto et al., 2006; Guzman et al., 2020). Leaf pieces were then floated on water (control) and suspensions (1x10^8^ CFU/mL) of *Pst* DC3000 or *Pf* A506. At the indicated times, leaf pieces were removed, dried with a filter paper, and placed on an Elite HD+ silicon rubber mixture (Zhermack) to create an impression of the leaf abaxial surface (Weyers and Johansen, 1985). Once hardened, the silicone rubber mixture was covered with clear nail varnish, which was allowed to dry, transferred to a glass slide, and observed under an Axio Zoom.V16 microscope (Zeiss). For each data point, the width and length of approximately 100-200 stomatal apertures were measured by using the Fiji package (https://fiji.sc/) and used to calculate the Stomatal Aperture Index. Images were also used to calculate the Stomata Number Index (Zhang et al., 2020).

### RNA-seq cDNA library preparation

Total RNA was isolated with the RNeasy Plant Mini Kit (QIAGEN) from wild-type and *bsk830-1* tomato leaves inoculated by vacuum with *P. fluorescens* A506 or mock-inoculated (three biological replicates for treatment, in total 12 samples collected at 6 h post-inoculation). RNA integrity was evaluated using the 4200 TapeStation System (Agilent Technologies). RNA-Seq cDNA libraries were prepared from RNA samples using the NEBNext Ultra^™^ II mRNA Library Prep Kit for Illumina and then PCR-amplified using NEBNext Multiplex Oligos for Illumina (New England Biolabs). Quality and average size of cDNAs in the library were evaluated using the 4200 TapeStation System (Agilent Technologies).

### Next-generation sequencing and data analysis

Libraries were sequenced using a NextSeq^™^ 500 system (Illumina) at the Genomics Research Unit of the Faculty of Life Sciences, Tel Aviv University. FASTQ files obtained from the sequencing were analyzed by Partek® Flow® 8.0 using the *Solanum lycopersicum* SL3.0 assembly (NCBI ID 393272). Raw reads with Phred quality scores of less than 20 were trimmed from the 3′ end, followed by removal of adaptor sequences. High quality trimmed reads (Phred score ∼34 and read length 75 bp) were aligned to the reference genome by STAR 2.7.3a (Dobin et al., 2013).

Gene expression quantification was performed using the Partek E/M algorithm (Xing et al., 2006), and normalized to RPKM (reads per kilobase of transcript per million mapped reads) (Mortazavi et al., 2008). Gene-specific analysis was performed on 15,575 detected genes (false discovery rate [pFDR] < 0.05). Overlap of differentially expressed genes in wild-type and *bsk830-1* plants was visualized using the eulerr R package (Larsson, 2020). Gene ontology (GO) enrichment analysis was performed using g:Profiler (version e105_eg52_p16_e84549f) with a significance threshold of pFDR < 0.05 (Raudvere et al., 2019). The term size of functional categories was limited to 10-150 to exclude GO terms associated with general processes, and electronic GO annotations were discarded to increase GO term accuracy.

### Statistical analysis

Experiments were performed at least three times. Statistical significance is based on either one- or two-way ANOVA followed by Tukey’s post-hoc test performed in the R environment. The multcompView R package (Graves et al., 2019) was used to assign a compact letter display to indicate the statistical differences in post-hoc tests.

## Accession numbers

Sequence data from this article can be found in the Solanaceae Genomics Network database (https://solgenomics.net/) under the following accession numbers: *Bsk880* (Solyc01g080880), *Bsk260* (Solyc04g082260), *Bsk600* (Solyc06g076600), *Bsk750* (Solyc09g011750), *Bsk000* (Solyc10g085000), *Bsk890* (Solyc11g064890), *Bsk830* (Solyc12g099830), *Fls2.1* (Solyc02g070890), *Fls2.2* (Solyc02g070910), *Fls3* (Solyc04g009640). RNA-seq reads generated in this work are available under accession number GSE199518 in the NCBI Gene Expression Omnibus (GEO) database.

## AKNOWLEDGMENTS

We thank Mary Beth Mudgett for providing the *Pst* DC3118 strain and Doron Teper for critical reading of the manuscript. This work was supported by grants from the United States-Israel Binational Agricultural Research and Development Fund (BARD; grant no. IS-4931-16C and IS-3541-21C to G.S. and G.B.M.) and the National Science Foundation (grant no. IOS-1546625 to G.B.M).

## AUTHOR CONTRIBUTIONS

G.So., B.B.M., G.B.M., and G.S. conceived and designed the experimental plans and analyzed the data; B.B.M. performed the protein-protein interaction analyses, and G.So. performed all the other experiments; N.Z. and H.M.R. generated the *bsk830* mutant plants; M.P.C. analyzed the RNA-seq data; G.So. and G.S. wrote the article.

## SUPPLEMENTAL TABLES AND FIGURE LEGENDS

**Supplemental Table S1.** List of differentially expressed genes.

**Supplemental Table S2.** Primers used in this study.

**Supplemental Figure S1.** A, Expression in yeast of the kinase domain of Fls2 (Fls2_KD_) and Fls3 (Fls3_KD_) fused to the GAL4 DNA-activation domain (GAL4-AD). B, Expression in leaves of *N. benthamiana* plants of Fls2_KD_ and Fls3_KD_ fused to the C-terminal half (C-LUC) of the luciferase protein. Proteins were detected by immunoblot analysis using anti-HA antibodies (α-HA) or anti-luciferase antibodies (α- LUC).

**Supplemental Figure S2.** Bsk830 is not phosphorylated *in vitro* by Fls2 and Fls3. Phosphorylation of the maltose binding protein (MBP)-Bsk830 fusion by the cytoplasmic domain of Fls2 (Fls2_CD_) and Fls3 (Fls3_CD_) fused to MBP was assayed *in vitro* in the presence of [γ-^32^P]ATP. Proteins were fractionated by SDS-PAGE, blotted onto a PVDF membrane and exposed to autoradiography, or stained with Coomassie Blue.

**Supplemental Figure S3.** Sequence of the *Bsk830* deletions in the tomato *bsk830-1* and *bsk830-2* mutant lines. Multiple sequence alignment of the *Bsk830* region flanking the gRNA-targeted site in wild-type, *bsk830-1*, and *bsk830-2* plants. In green, the ATG translation start site; in blue, the PAM motif; in pink, the gRNA. A red dotted line represents sequences deleted in the mutant lines.

**Supplemental Figure S4.** Tomato *bsk830* mutant plants are not impaired in flg22- and flgII-28-induced MAPK activation. Leaf discs of wild-type, *bsk830-1, bsk830-2, fls2.1/fls2.2* and *fls3* mutant plants were floated overnight in water and treated with 1 μM of flg22 (A) or flgII-28 (B). Samples were harvested at 0, 5 and 20 min after treatment and analyzed by immunoblots with anti-pMAPK antibodies (α-pMAPK). Ponceau S staining of RuBisCO is shown as a loading control. Data are representative of three biological repeats.

**Supplemental Figure S5.** Stomatal number index of wild-type, *bsk830-1,* and *bsk830-2* tomato plants. The number of stomata and epidermal pavement cells was manually counted in a 0.5 mm^2^ leaf area and the stomatal number index was calculated as the percentage of stomata per total cells. Approximately 30 images were analyzed for each plant genotype.

