## Supplemental Figure S for "Tomato brassinosteroid-signaling kinase Bsk830 is a component of flagellin signaling that regulates pre-invasion immunity"

### Sobol et al., Supplemental Figure S1

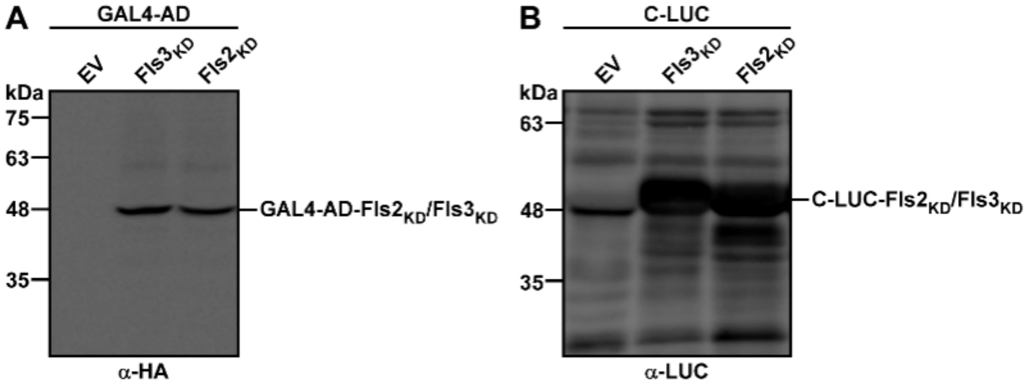


**Supplemental Figure S1.** A, Expression in yeast of the kinase domain of Fls2 ($Fls2_{KD}$) and Fls3 ($Fls3_{KD}$) fused to the GAL4 DNA-activation domain (GAL4-AD). B, Expression in leaves of *N. benthamiana* plants of $Fls2_{KD}$ and $Fls3_{KD}$ fused to the C-terminal half (C-LUC) of the luciferase protein. Proteins were detected by immunoblot analysis using anti-HA antibodies (α-HA) or anti-luciferase antibodies (α-LUC).

**Sobol et al., Supplemental Figure S2**

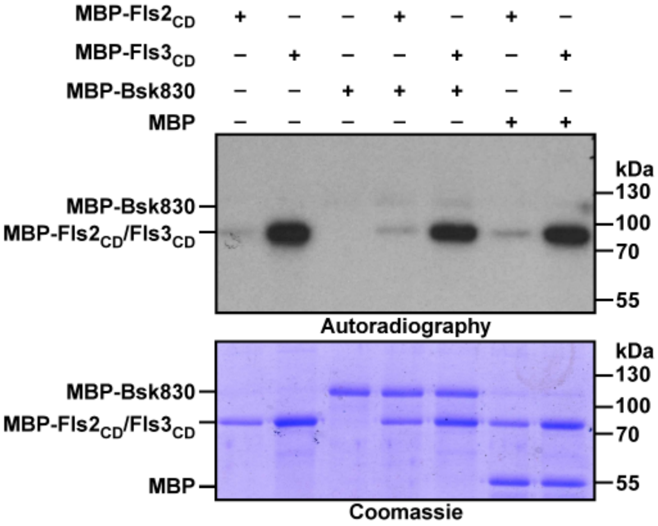


**Supplemental Figure S2.** Bsk830 is not phosphorylated *in vitro* by Fls2 and Fls3. Phosphorylation of the maltose binding protein (MBP)-Bsk830 fusion by the cytoplasmic domain of Fls2 ($Fls2_{CD}$) and Fls3 ($Fls3_{CD}$) fused to MBP was assayed *in vitro* in the presence of [$\gamma$-$^{32}$P]ATP. Proteins were fractionated by SDS-PAGE, blotted onto a PVDF membrane and exposed to autoradiography, or stained with Coomassie Blue.

### Sobol et al., Supplemental Figure S3

```
                                                                    PAM        gRNA1
Wild-type  ATGGGCTGTGAAAGTTCTAAACTTGCCTCATGTTGCTGGACTGGAGAGAGTGGCCCCATTCACGAGGCTCAGAATCC   77
bsk830-1   ATGGGCTGTGAAAGTTCTAAACTTGCCTCATGTTGCTGGACTGGAGAGAGTGGCCCCAT----GAGGCTCAGAATCC
bsk830-2   AT---------------------------------------------------------------------------

Wild-type  TGGTATGTATATCTTTTATTGAAATATTTTCTATATAACCATCCTACTGCAGTGTTTAATAAATTTTTTTTCACGCT  154
bsk830-1   TGGTATGTATATCTTTTATTGAAATATTTTCTATATAACCATCCTACTGCAGTGTTTAATAAATTTTTTTTCACGCT
bsk830-2   ------------------------------------------------------------ACTATATCTTTCACGCT

Wild-type  TCTCGTAGATGAAGAAAAAAAATGAAGTTAGTGATTCACCTGCATTCTGCGAGTTTACATTTGAGCAGCTCAGGATAG  231
bsk830-1   TCTCGTAGATGAAGAAAAAAAATGAAGTTAGTGATTCACCTGCATTCTGCGAGTTTACATTTGAGCAGCTCAGGATAG
bsk830-2   TCTCGTAGATGAAGAAAAAAAATGAAGTTAGTGATTCACCTGCATTCTGCGAGTTTACATTTGAGCAGCTCAGGATAG

Wild-type  CTACATCTGCATTTGCTGTAGAGAATATAGTTTCTGAACATGGCGAAAAAGCTCCAAATGTTGTTTATAAAGGAAAG  308
bsk830-1   CTACATCTGCATTTGCTGTAGAGAATATAGTTTCTGAACATGGCGAAAAAGCTCCAAATGTTGTTTATAAAGGAAAG
bsk830-2   CTACATCTGCATTTGCTGTAGAGAATATAGTTTCTGAACATGGCGAAAAAGCTCCAAATGTTGTTTATAAAGGAAAG

                   PAM        gRNA2
Wild-type  CTAGAGAACCAGAGGCCGGTTGCTGTTAAACGCTTCAACAGATCTGCATGGCCTGATTCCCGGCAATTTTTAGTAAG  385
bsk830-1   CTAGAGAACCAGAGGCCGGTTGCTGTTAAACGCTTCAACAGATCTGCATGGCCTGATTCCCGGCAATTTTTAGTAAG
bsk830-2   CTAGAGAACCAGAGGCCGGTTGCTGTTAAACGCTTCAACAGATCTGCATGGCCTGATTCCCGGCAATTTTTAGTAAG
```

**Supplemental Figure S3.** Sequence of the *Bsk830* deletions in the tomato *bsk830-1* and *bsk830-2* mutant lines. Multiple sequence alignment of the *Bsk830* region flanking the gRNA-targeted site in wild-type, *bsk830-1*, and *bsk830-2* plants. In green, the ATG translation start site; in blue, the PAM motif; in pink, the gRNA. A red dotted line represents sequences deleted in the mutant lines.

### Sobol et al., Supplemental Figure S4

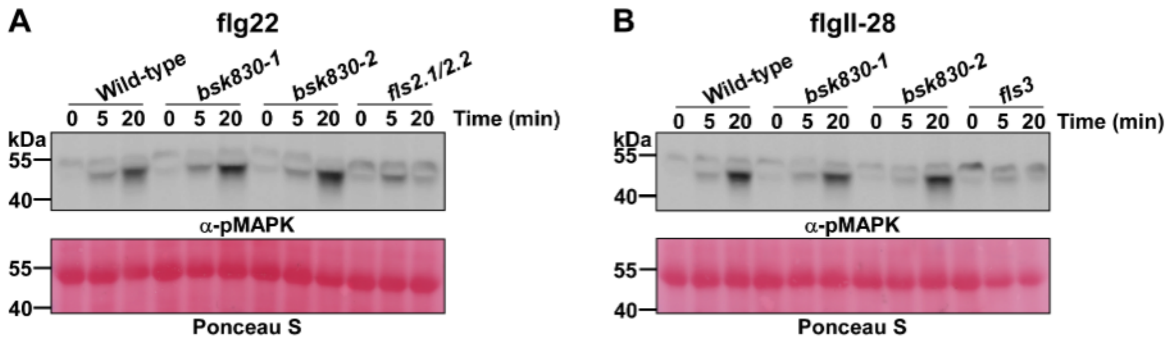


**Supplemental Figure S4.** Tomato *bsk830* mutant plants are not impaired in flg22- and flgII-28-induced MAPK activation. Leaf discs of wild-type, *bsk830-1*, *bsk830-2*, *fls2.1*/*fls2.2* and *fls3* mutant plants were floated overnight in water and treated with 1 μM of flg22 (A) or flgII-28 (B). Samples were harvested at 0, 5 and 20 min after treatment and analyzed by immunoblots with anti-pMAPK antibodies (α-pMAPK). Ponceau S staining of RuBisCO is shown as a loading control. Data are representative of three biological repeats.

### Sobol et al., Supplemental Figure S5

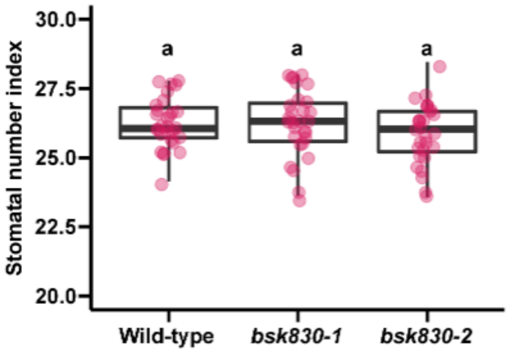


**Supplemental Figure S5.** Stomatal number index of wild-type, *bsk830-1*, and *bsk830-2* tomato plants. The number of stomata and epidermal pavement cells was manually counted in a 0.5 $mm^2$ leaf area and the stomatal number index was calculated as the percentage of stomata per total cells. Approximately 30 images were analyzed for each plant genotype.
