## Supplemental Table S2 for "Tomato brassinosteroid-signaling kinase Bsk830 is a component of flagellin signaling that regulates pre-invasion immunity"

**Supplemental Table S2.** Primers used in this study.

| **Primer ID** | **Name** | **Sequence (5’ to 3’)** | **Purpose** |
| --- | --- | --- | --- |
| 1 | FLS3KD_F_SfiI | AGTGGCCATTACGGCCCATGTTTGATGAATCCAATTTGATTGG | Cloning of Fls3_KD_ into pGADT7-RecM for Y2H |
| 2 | FLS3KD_R_SfiI | AAAGGCCGAGGCGGCCCTAATTTACTTCTATGTTTC |  |
| 3 | FLS2KD_F_SfiI | AGTGGCCATTACGGCCCATGCATGCTACCAATAATTTCCGTCCGG | Cloning of Fls2_KD_ into pGADT7-RecM for Y2H |
| 4 | FLS2KD_R_SfiI | AGAGGCCGAGGCGGCCATTAGCCACCATGGGAAACTTGGT |  |
| 5 | FLS3KD_F_KpnI | ACGGGGTACCATGTTTGATGAATCCAATTTGATTGG | Cloning of Fls3_KD_ into pCAMBIA:C-LUC for SLCA |
| 6 | FLS3KD_R_SalI | ACGCGTCGACCTAATTTACTTCTATGTTTC |  |
| 7 | FLS2KD_F_KpnI | ACGGGGTACCATGCATGCTACCAATAATTTCCGTCCG | Cloning of Fls2_KD_ into pCAMBIA:C-LUC for SLCA |
| 8 | FLS2KD_R_SalI | ACGCGTCGACTTAATCTTTTACCAAATGAGAAGGC |  |
| 9 | BSK830_F_KpnI | TTTTTTGGTACCATGGGCTGTGAAAGTTCTAAACTTGCCTC | Cloning of Bsk830 into pBTEX:YFP |
| 10 | BSK830_R_XbaI | TTTTTTTCTAGAAGCAGATGCATTTTTCTCTTCTTC |  |
| 11 | BSK830_G2A_F_KpnI | TTTTTTGGTACCATGGCCTGTGAAAGTTCTAAACTTGCCTC | Cloning of Bsk830^G2A^ into pBTEX:YFP |
| 12 | BSK830_C3,11,12A_F_KpnI | TTTTTTGGTACCATGGGCGCTGAAAGTTCTAAACTTGCCTCAGCTGCCTGGACTGGAGAGAG | Cloning of Bsk830^C(3,11,12)A^ into pBTEX:YFP |
| 13 | BSK830_F_BamHI | ACGGGGATCCATGGGCTGTGAAAGTTCTAAACTTGC | Cloning of Bsk830 into pMAL-C2X |
| 14 | BSK830_R_SalI | ACGCGTCGACTCAAGCAGATGCATTTTTCTCTTCTTCG |  |
| 15 | BSK830_F_sgRNA_flanking | CGACTCCAACTAGTTTATGTTCAAGG | Verification of mutation site in *Bsk830* |
| 16 | BSK830_R_sgRNA_flanking | GTTTCCATTTTGTTGTTGATGCATTAGG |  |
